# Thermal preference influences depth use but not biomass of predatory fishes in response to lake morphometry

**DOI:** 10.1101/572925

**Authors:** Timothy J. Bartley, Matthew M. Guzzo, Kevin Cazelles, Alex Verville, Bailey C. McMeans, Kevin S. McCann

## Abstract

Top predators’ responses to environmental conditions shape food web architecture and influence ecosystem structure and stability. Yet the impacts of fundamental properties like ecosystem size and morphometry on top predators’ behaviour are poorly understood. We examined how lake morphometry impacts the behaviour (inferred by depth use) of three key fish top predators—the cold-adapted lake trout, the cool-adapted walleye, and the warm-adapted smallmouth bass— which can each strongly impact local food web structure. We used catch-per-unit-effort data from nearly 500 boreal lakes of Ontario, Canada to evaluate the role of thermal preference in dictating mean depth of capture and biomass index in response to lake morphometry. We found evidence that thermal preferences influence how species’ depth use and biomass changed with lake size, proportion of littoral area, and maximum lake depth, although we found no relationship with lake shape. However, found no strong evidence that lake morphology influences these species’ biomasses, despite theory that predicts such a relationship. Our results suggest that some aspects of lake morphometry can alter habitat accessibility and productivity in ways that influence the behaviour and biomass of these top predator species depending on their thermal preferences. Our results have implications for how lake food webs expand and contract with lake morphometry and other key abiotic factors. We argue that several key abiotic factors likely drive top predator depth use in ways that may shape local food web structure and play an important role in determining the ultimate fate of ecosystems with environmental change.

## INTRODUCTION

Ecosystem size and shape have long intrigued ecologists (Elton 1927) and are widely acknowledged as fundamentally important in ecology. As a result, ecologists have developed rich theory about how these factors govern species diversity, identity, and interactions in an ecosystem (MacArthur and Wilson 1967, Post et al. 2000, Tunney et al. 2012). These morphometric properties are often accounted for in ecological research; however, they are less often the primary focus in the ecology literature despite their potential to filter the effects of global environmental change on ecosystems (MacArthur and Wilson 1967, Jackson et al. 2001, Fahrig 2003). Understanding the role of ecosystem size and shape alone and in combination with other environmental factors is paramount in this time of rapid global environmental change.

Top predators are often large, mobile, and opportunistic, allow them to behaviourally respond to environmental conditions in ways that shape their fitness and their supporting food webs (Cruz-Font et al. 2019, Hammerschlag et al. 2019). One such behaviour is habitat coupling, where predators can move and feed between spatially distinct macrohabitats (Schindler and Scheuerell 2002). Habitat coupling appears to be a fundamental component of food architecture that, importantly, can stabilize food webs (Rooney et al. 2008, McCann and Rooney 2009). Consumers can rapidly switch away from low-density resources and towards other, more abundant resources, creating a stable resource base from the perspective of consumers that produces relatively stable consumer densities. This switching also simultaneously culls high-density prey and relieves low-density prey from top-down pressure (McCann and Rooney 2009, Valdovinos et al. 2010).

Top predators can influence the trophic biomass structure of ecosystems through their coupling of habitats. In theory, when consumers couple distinct habitats, they have access to a larger overall energy pool, which ought to inflate in their biomass (McCann et al. 2005, McCauley et al. 2018). These changes in biomass can, in turn, affect a predator’s ability to suppress prey through top-down control, leading to top-heavy food webs and impacting stability (McCann et al. 2005, McCauley et al. 2018). This theoretical prediction has some empirical support. For example, increased habitat coupling is associated with increased biomass of the predatory walleye (*Sander vitreus*) (Tunney et al. 2018). Yet, widespread empirical evidence is lacking for whether the shifts in habitat coupling exhibited by predators translate to altered predator biomasses.

The degree to which predatory fish couple offshore and nearshore habitats is potentially shaped by a few key factors. One possibility is that predators respond to changes in resource abundance by shifting towards locations with abundant resources (i.e., the birdfeeder effect, see Eveleigh et al. 2007). This phenomenon has been documented in the freshwater top predator northern pike (*Esox lucius*) (Kennedy et al. 2018). However, because fish are ectotherms, the ability of predatory fish to respond to resource abundances may be limited by temperature (Tunney et al. 2012, 2014, Guzzo et al. 2017). The thermal preferences of fishes generally fall into three guilds—cold-water, cool-water and warm-water (Magnuson et al. 1979). During summer, lakes thermally stratify, and warm surface waters often exceed the temperature preferences of cold-water fish, increasing the metabolic costs associated with occupying shallow habitats. In response, the cold-adapted predatory fishes respond by retreating to the deep water and at times accessing nearshore resources in quick forays (Bertolo et al. 2011, Guzzo et al. 2017, Cruz-Font et al. 2019). Cold-water fishes also exhibit similar responses across natural gradients in lake temperature (Tunney et al. 2014, Guzzo et al. 2017, Bartley et al. 2019), suggesting that thermal accessibility may be limiting for many species. Various morphometric factors appear to influence the accessibility of the nearshore in lakes. During summer, large and reticulate lakes likely have larger nearshore zones, and predatory fish must spend more time and travel greater distances through warm water to reach prey (Dolson et al. 2009). As a consequence, increasing lake size and complexity are associated with reduced consumption less from littoral resource pathways in lake trout (Dolson et al. 2009, Tunney et al. 2012). Importantly, these foraging shifts are associated with behavioural shifts in the form of altered habitat use (Guzzo et al. 2017, Bartley et al. 2018). However, cool-water species such as the walleye are likely less thermally restricted than cold-water species, and so ought to show weak behavioural response to changes in nearshore accessibility (Fig. 1). Furthermore, warm-water species, such as smallmouth bass, may never be thermally restricted from accessing prey in the nearshore and thus may show no behavioural responses to lake morphometry (Fig. 1).

**Figure 1.**
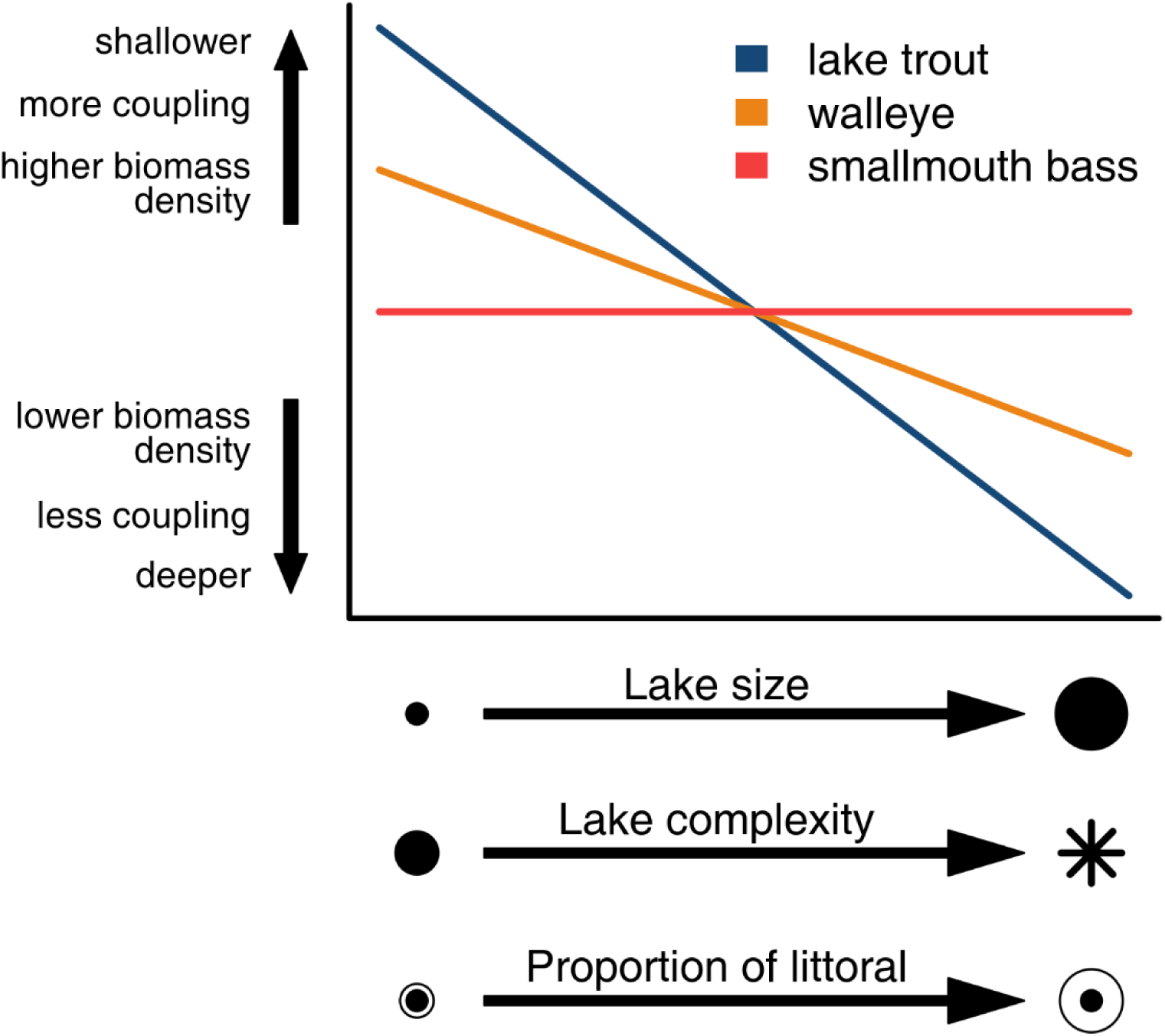
Predictions of the influence of lake surface area and shape (i.e., shoreline development index) on the habitat use and biomass of three top predator species from three thermal guilds: the cold-adapted lake trout (*Salvelinus namaycush*), the cool-adapted walleye (*Sander vitreus*), and the warm-adapted smallmouth bass (*Micropterus dolomieu*).

Changes in habitat coupling by top predators likely have important consequences for lake food webs (McCann et al. 2005, Hammerschlag et al. 2019). If habitat coupling does indeed increase the overall resource pool available to predators, inflating predators’ biomasses (McCann et al. 2005, McCauley et al. 2018), factors like lake morphometry may impact the ability fish predators to suppress their prey and respond to changes in resource density. There is some empirical evidence that predators’ biomass responds to changes in lake size (Tunney et al. 2012, 2018, Eloranta et al. 2015). Thus, the differences in coupling responses to changes in lake morphometry predicted here ought to produce a large reduction in energy availability for lake trout but little reduction for walleye and no reduction for smallmouth bass. Thus, these three species ought to show contrasting biomass responses to changes in lake size and complexity. Such responses would strongly suggest that lake morphometry shapes whole food webs.

Here, we use an extensive catch-per-unit-effort database of fishes in over nearly 500 boreal lakes of Ontario, Canada to evaluate the role of thermal preference in driving the depth use (inferred from mean depth of capture) and biomass index responses to lake morphometry. We predict that as lake size, complexity and proportion littoral area increase, lake trout will be caught at much deeper depths (Fig. 1). We predict that walleye will show weak depth response to lake morphometry, and that smallmouth bass will show no depth response to changes in lake morphometry (Fig. 1). Because changes in depth use by fish are thought to influence species biomass through changes in resource availability, we also predict that lake trout will show decreasing biomass index with increasing lake size, complexity, and proportion littoral area. We predict that walleye will show weak biomass index decreases with these lake morphometric variables, and that smallmouth bass will show no change in biomass index. By evaluating the depth use and biomass of these three key top predator species with changes in habitat accessibility, we provide further insight into the response of mobile generalist consumers to key ecosystem characteristics that can shape the ways that ecosystems are altered by global environmental change.

## METHODS

### Data Source

We used an extensive catch-per-unit-effort (CPUE) database of 721 lakes constructed by the Ontario Ministry of Natural Resources and Forestry (OMNRF) Broad-scale Fisheries Monitoring (BsM) Program (Sandstrom et al. 2013). The BsM program includes long-term monitoring of fish populations and communities, physical and chemical water characteristics, aquatic habitat, invasive species, and fishing effort. BsM fish surveys are designed to quantitatively assess the distribution and abundance of fish species and characterize the diversity and identity of fish communities throughout Ontario. Fish surveys take place during one period in summer (typically between May and September, when water surface temperatures are at least 18°C). BsM surveys are conducted using a combination of two types of benthic gillnets: large mesh nets (North American standard multipaneled gill nets with mesh sizes varying from 38 to 127 mm) and small-mesh nets (mesh sizes from 13 to 38 mm). Gillnets are set overnight, perpendicular to shore and randomly throughout the extent of each lake, with the number of nets set determined by lake size and depth (for details, see Sandstrom et al. 2013). Large mesh nets are set in each of eight depth strata (1-3m, 3-6m, 6-12m, 12-20m, 20-35m, 35-50m, 50-75m, and >75m) that is present in at least 5% of the surface area of the lake. Small mesh nets are also stratified but set only in the first four of these depth strata. More information about the BsM sampling program can be found through the Ontario Ministry of Natural Resources and Forestry website (https://www.ontario.ca/page/broad-scale-monitoring-program) and in Sandstrom et al. 2013.

For our study, we used data from Cycle 1 of BsM, which took place from 2008 to 2012. We only included lakes with a maximum depth greater than 10 m to ensure that both nearshore and offshore habitats were present for the lakes that we use in our analysis. This criterion left 556 lakes for our analyses. Because the BsM protocol uses a relatively small number of fixed depth strata (especially in the littoral zone), it is limited in its ability to detect depth use change within the nearshore habitat but is well suited to detect changes in depth use in terms of nearshore and offshore habitat use.

### Lake Attribute Selection

We selected seven lake attributes to use as predictor variables in our statistical analysis following a two-step procedure. First, we retrieved 10 lake attributes available for the 556 lakes included in our analysis from OMNRF’s data to evaluate lake size, lake complexity, and habitat availability, but also variables to control for climatic conditions, and productivity. Lake size was measured by the surface area of the lake and lake complexity by the shoreline development index (or SDI) computed following Dolson et al. (2009) using lake area and lake shoreline. We used growing degree days based on air temperature (yearly average cumulative number of degrees above 5°C from 1981 to 2010) and latitude as proxies for climatic condition; Secchi depth, total phosphorus and dissolved organic carbon to assess productivity; mean and maximum depth (based on the bathymetry of the lake), and the proportion of littoral area (proportion of the lake less than 4.6 m) to assess habitat availability. In a second step, we transformed all these variables (see Table 1) and scaled them in order to perform a principal component analysis (PCA). Based on the results (see Fig S1, Table S1 and S2), we narrowed down the control variables to two: growing degree days for climatic conditions and Secchi depth for productivity. Because of strong correlations between mean depth and maximum depth (r = 0.85) and between mean depth and proportion littoral area (0.89) but relatively weaker correlation between maximum depth and proportion littoral area (0.65), we excluded mean depth from our analysis and included both maximum depth and proportion littoral area. Because prey availability may also influence the responses of these predator species, we also investigated how the proportion of prey in the offshore changes with our abiotic variables.

**Table 1.** Summary of lake characteristics. The mean, median, standard deviation (SD), minimum and maximum for each of the abiotic variables used in our analyses as well as the transformation applied to normalize each variable for statistical analysis.

| Variable | Mean | Median | SD | Range | Transformation |
| --- | --- | --- | --- | --- | --- |
| surface area (ha) | 2555 | 978 | 6911 | 41–90484 | log |
| maximum depth (m) | 35.5 | 28.0 | 24.4 | 10.0–213.5 | log |
| mean depth (m) | 10.2 | 8.1 | 6.7 | 1.4–40.1 | log |
| shoreline development index (unitless) | 5.3 | 4.3 | 3.4 | 1.3–20.9 | log |
| proportion littoral area (unitless) | 0.39 | 0.37 | 0.19 | 0.06–0.99 | logit |
| growing degree days (°C) | 1630 | 1626 | 202 | 1059–2246 | none |
| Secchi depth (m) | 4.1 | 3.9 | 2.0 | 0.1–13.4 | log |

### Species Selection and Characteristics

We selected one top predator species from each of the three thermal guilds for Canadian freshwater fishes that are well distributed across lakes of Ontario. Lake trout (*Salvelinus namaycush*) is a cold-water top predator of offshore habitats in boreal shield lakes (Hasnain et al. 2013), and much research on behavioural responses to thermal accessibility has focused on this species (Morbey et al. 2006, Dolson et al. 2009, Plumb and Blanchfield 2009, Tunney et al. 2014, Guzzo et al. 2017, Bartley et al. 2019). Walleye (*Sander vitreus*) is a cool-adapted voracious piscivore and popular sport fish that is present in lakes across the boreal shield (Hasnain et al. 2013). Smallmouth bass (*Micropterus dolomieu*) is a warm-water adapted generalist top predator that is non-native in part of its range in Ontario (Hasnain et al. 2013, Alofs and Jackson 2015). These three species—all of which are popular sport fish for anglers (Queen’s Printer for Ontario 2015)—are among the most common species in Ontario and comprise a large portion of the average catch in the BsM database. Although the BsM survey data does not account for any differences in the catchability of these different species, our analyses compare the behaviour and catch within a species using scaled data, and thus differences in species’ catchability should not affect our results.

### Catch-per-unit-effort based Behavioural Metric

For our analysis, we used catch-per-unit-effort data to calculate a weighted mean depth of capture for each species, an approach that has been used in previous studies (Kennedy et al. 2018, Bartley et al. 2019). We used species-specific catch-per-unit-effort data converted into the number of individuals per 100 metres per night (CUE, provided by OMNRF) to make catch data comparable within and across lakes. For our analysis for each species, we examined lakes that had at least 10 individuals caught to ensure that our behavioural metrics were representative of the species depth distribution in the lake. This leaves 485 lakes with at least one of our three focal species for our analyses. Following previous studies (Bartley 2017, Kennedy et al. 2018, Bartley et al. 2019), we used BsM catch data for each depth stratum in each lake to examine the mean depth of capture for each species, calculated as

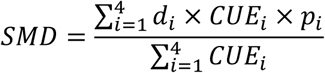

where SMD is the mean depth of capture of a fish species, *CUE*_*i*_ is the CUE of that species for depth stratum *i*, *p*_*i*_ is the proportion of the lake in depth stratum *i*, and *d*_*i*_ is the middle depth of the depth range for that depth stratum.

The stratified sampling protocol use for BsM means that the survey of each lake is essentially a “snapshot” of the habitat use (or depth distribution) of each species. We are using this snapshot to calculate the mean “behaviour” of a species—a value that typifies what that species is doing on average in that lake. We interpret shifts in the averages of this metric of average habitat use across lakes as inference the behavioural response of species to changing temperature. This approach of looking at changes in some metric across a gradient is common using other measures, such as stable isotope signatures (Dolson et al. 2009, Tunney et al. 2014). In addition, because only two depth strata were sampled in water < 6 meters during BsM surveys, our mean depth of capture metric likely does not capture changes in habitat use within the nearshore. For this reason, we included only lakes with a maximum depth of at least 10 m for our analysis. In these deeper lakes, enough strata were sampled to capture the relative use of offshore and nearshore habitats.

### Biomass Index

As an index of biomass we calculated the biomass per unit effort (WUE) for each species in each lake, we transformed the mean fork length of each species in each lake into a mean weight using the following equations: For lake trout, weight = 10^−5.19^ * (fork length)^3.1^; for walleye, weight = 10^−4.79^ * (fork length)^2.9^; for smallmouth bass, weight = 10^−4.7^ * (fork length)^3^. We then multiplied the CUE for each species in each lake by the estimated mean weight to get a mean weight per unit effort in kg per 100 metres per night or WUE, hereafter referred to as ‘biomass index’. Both fork length and CUE data as well as the equations we used to calculate WUE were provided by the Ontario Ministry of Natural Resources and Forestry.

### Prey fish distribution

We also calculated the proportion of prey fish captured in the offshore zone of each lake. We used the total number of prey fish captured in offshore nets (>6 m depth) divided by the total number of prey fish caught in that lake. We considered a species to be a prey fish if it was not primarily piscivorous in inland lakes (Coker et al. 2001). Our calculations included data from both small mesh and large mesh nets, but we only included species with at least 10 individuals <250 mm in length for our calculation to remove rare species and individuals too large to be prey for most predators. These criteria left 53 species as prey fish.

### Statistical Analyses

We used multiple regression to test the influence of physical lake characteristics on the depth of capture (a measure of depth use) and biomass index of predatory fish (lake trout, walleye, and smallmouth bass) as well as the proportion of prey fish captured in the offshore of lakes, which may affect the depth use of these predator species. For each response variable, model selection was performed using backwards selection based on AIC starting with a starting model that contained the seven predictor variables without interactions (Table 1). The predictor variables were transformed to meet the assumptions of multiple regression (Zuur et al. 2010).

Because BsM surveys sampled every depth stratum present in a given lake, the number of depth strata sampled in each lake increases with lake depth. To determine whether our results were driven by the presence of more, deeper depth strata with increasing lake depth, we performed our regression analysis on the subset of lakes that were sampled in each of the first 5 depth strata (from 1 to 35 m) but not in depth strata 6 – 8 (greater than 35 m). Each lake in this subset has at least 5% of surface area with depths of 20-35m and no more than 5% of surface area greater than 35 m. Because the results of regressions performed on this subset are similar to the results using all lakes (see Figs. S2, S3, and S4), we are confident that our results here are not simply the result of the sampling methodology.

To better understand how our results for each species’ biomass index were being independently influenced by body size and CUE, we also performed our regressions separately on the two variables used to calculate biomass index: catch-per-unit-effort and mean weight (Figs. S6 and S7).

## RESULTS

### Depth of capture

For lake trout, the top-ranked model accounted for much of the variation in mean depth of capture among lakes (57%) and contained the predictor variables lake surface area, lake maximum depth, growing degree days, proportion littoral area, and shoreline development index (Table 2). Scaled coefficients indicated that as lake surface area, lake maximum depth, and growing degree days increased, lake trout were captured at deeper depths, with lake surface area having the strongest influence of these three variables (Table 2, Fig. 2, 3). Proportion littoral area and shoreline development index each had relative weak influences on lake trout depth (Table 2).

**Table 2.** Results of the stepwise model selection of multiple regression testing for the influence of physical lake characteristics on the mean depth of capture of lake trout, walleye, and smallmouth bass in study lakes. SE is standard error.

| Species | Variable | Scaled coefficient | SE | t-value | Significance<sup>1</sup> | df | F-value | R<sup>2</sup><sub>adj</sub> |
| --- | --- | --- | --- | --- | --- | --- | --- | --- |
| lake trout | intercept | 0.000 | 0.047 | 0.000 |  | 5, 184 | 51.81 | 0.573 |
|  | surface area | 0.453 | 0.080 | 5.663 | *** |  |  |  |
|  | shoreline development index | -0.128 | 0.073 | -1.762 | . |  |  |  |
|  | growing degree days | 0.173 | 0.049 | 3.561 | *** |  |  |  |
|  | maximum depth | 0.287 | 0.072 | 3.979 | *** |  |  |  |
|  | proportion littoral area | -0.186 | 0.066 | -2.820 | ** |  |  |  |
| walleye | intercept | 0.000 | 0.049 | 0.000 |  | 4, 320 | 26.053 | 0.236 |
|  | surface area | 0.316 | 0.055 | 5.756 | *** |  |  |  |
|  | growing degree days | -0.075 | 0.049 | -1.526 |  |  |  |  |
|  | maximum depth | -0.223 | 0.069 | -3.239 | ** |  |  |  |
|  | proportion littoral area | -0.521 | 0.064 | -8.193 | *** |  |  |  |
| smallmouth bass | intercept | 0.000 | 0.060 | 0.000 |  | 5, 206 | 13.493 | 0.228 |
|  | shoreline development index | 0.199 | 0.067 | 2.977 | ** |  |  |  |
|  | growing degree days | 0.286 | 0.065 | 4.370 | *** |  |  |  |
|  | maximum depth | -0.208 | 0.090 | -2.317 | * |  |  |  |
|  | Secchi depth | 0.134 | 0.068 | 1.977 | * |  |  |  |
|  | proportion littoral area | -0.528 | 0.092 | -5.728 | *** |  |  |  |

<sup>1</sup>P-value significance: 0 ‘\*\*\*’ 0.001 ‘\*\*’ 0.01 ‘\*’ 0.05 ‘.’ 0.1 ‘ ’ 1

**Figure 2.**
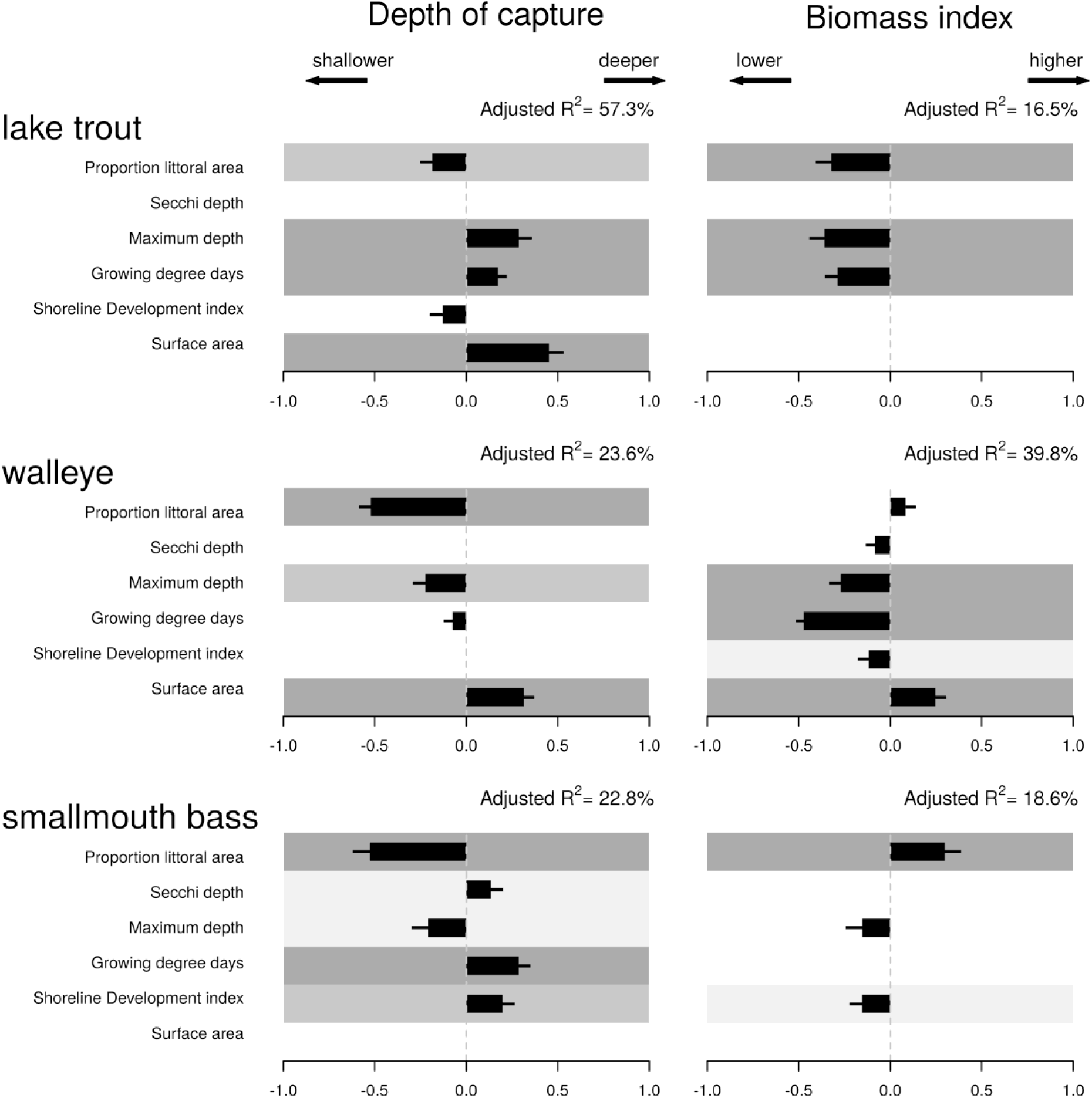
Impact of predictors on depth of capture (left side) and biomass index (right side) for lake trout, walleye, and smallmouth bass. The scaled effect of each variable included for a species and the corresponding standard error are depicted with black bars. A gray background is added to all explanatory variables found statistically significant (the different p-value thresholds 0.05, 0.01 and 0.001 are coloured in light, medium and dark gray, respectively).

**Figure 3.**
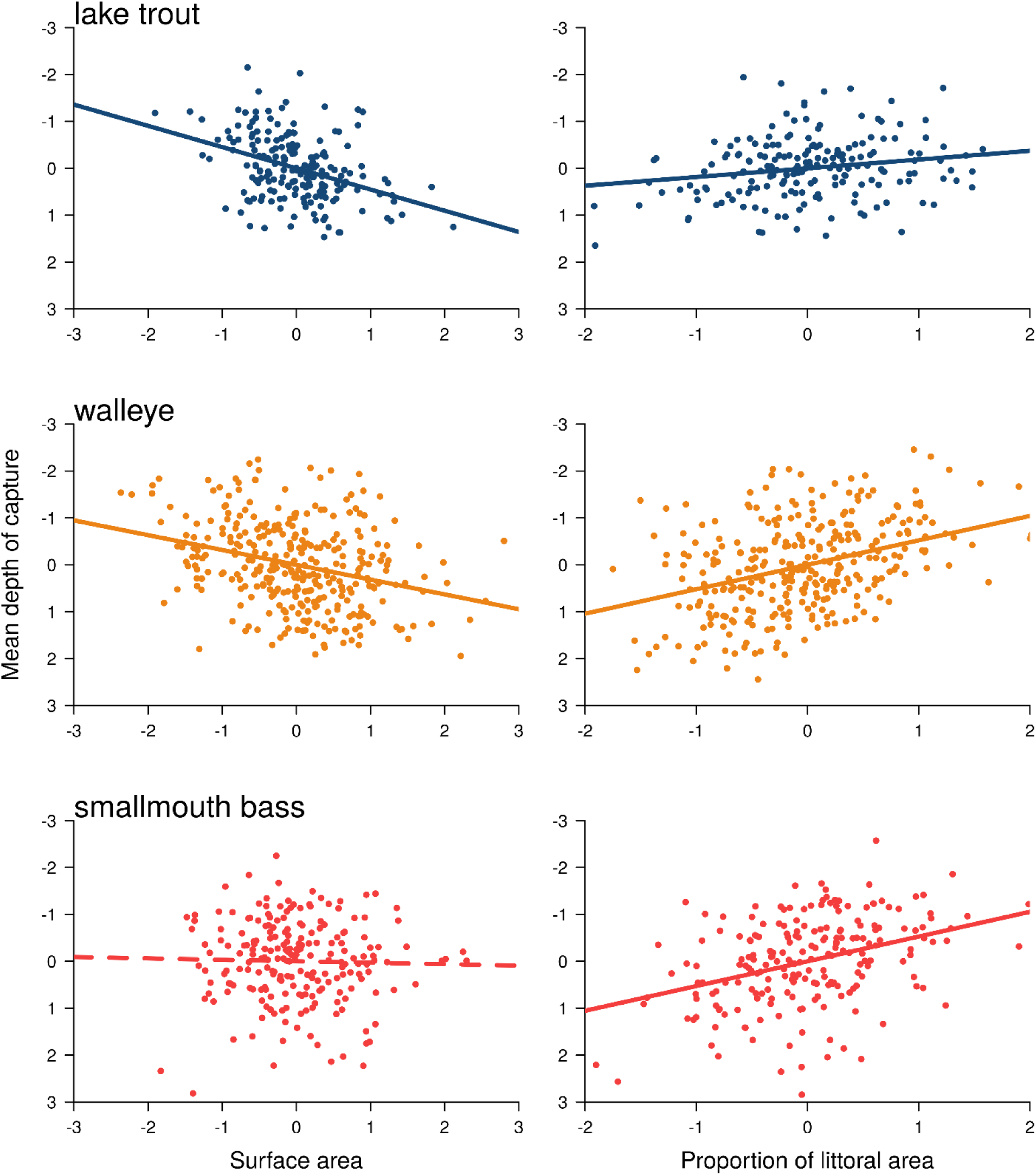
Added variable plots depicting relationships between mean depth of capture of lake trout, walleye, and smallmouth bass with lake surface area and proportion of littoral area in Ontario lakes.

For walleye, the top-ranked model accounted for 24% of the variation in mean depth of capture among lakes and contained the predictor variables lake surface area, maximum depth, growing degree days, and proportion of littoral area (Table 2). Scaled coefficients indicated that lake surface area had a positive effect on depth of capture, while the proportion of littoral area, maximum depth had strong and moderate negative effects on mean depth of capture, respectively (Table 2, Fig. 2, 3). Although growing degree days was included in the top-ranked model, this variable did not have an obvious influence on depth of capture (Table 2, Fig. 2).

For smallmouth bass, the best model accounted for 23% of the variation in mean depth of capture among lakes and contained the predictor variables lake maximum depth, growing degree days, proportion littoral area, Secchi depth, and shoreline development index (Table 2). Scaled coefficients indicated shoreline development index, growing degree days, and Secchi depth each had a relatively weak positive effect on mean depth of capture, while proportion of littoral area and lake depth had a strong and moderate negative effect on mean depth of capture, respectively (Table 2, Fig. 2, 3).

### Biomass Index

For lake trout, the top-ranked model accounted for 17% of the variation in biomass index among lakes and contained the predictor variables lake maximum depth, growing degree days, and proportion littoral area (Table 3). Scaled coefficients indicated that each of these variables had moderate negative effects on lake trout biomass index (Table 3, Fig. 2, 4).

**Table 3.** Results of the stepwise model selection of multiple regression testing for the influence of physical lake characteristics on the biomass index of lake trout, walleye, and smallmouth bass in study lakes. SE is standard error.

| Species | Variable | Scaled coefficient | SE | t-value | Significance<sup>1</sup> | df | F-value | R<sup>2</sup><sub>adj</sub> |
| --- | --- | --- | --- | --- | --- | --- | --- | --- |
| lake trout | intercept | 0.000 | 0.066 | 0.000 |  | 3, 186 | 13.445 | 0.165 |
|  | growing degree days | -0.288 | 0.067 | -4.264 | *** |  |  |  |
|  | maximum depth | -0.359 | 0.083 | -4.329 | *** |  |  |  |
|  | proportion littoral area | -0.323 | 0.083 | -3.883 | *** |  |  |  |
| walleye | intercept | 0.000 | 0.043 | 0.000 |  | 6, 318 | 36.762 | 0.398 |
|  | surface area | 0.246 | 0.061 | 4.026 | *** |  |  |  |
|  | shoreline development index | -0.118 | 0.058 | -2.031 | * |  |  |  |
|  | growing degree days | -0.472 | 0.045 | -10.490 | *** |  |  |  |
|  | maximum depth | -0.271 | 0.064 | -4.256 | *** |  |  |  |
|  | Secchi depth | -0.084 | 0.050 | -1.678 | . |  |  |  |
|  | proportion littoral area | 0.083 | 0.059 | 1.405 |  |  |  |  |
| smallmouth bass | intercept | 0.000 | 0.062 | 0.00 |  | 3, 208 | 17/039 | 0.186 |
|  | shoreline development index | -0.155 | 0.069 | -2.248 | . |  |  |  |
|  | maximum depth | -0.153 | 0.091 | -1.682 | . |  |  |  |
|  | proportion littoral area | 0.298 | 0.089 | 3.345 | *** |  |  |  |

<sup>1</sup>P-value significance: 0 '\*\*\*' 0.001 '\*\*' 0.01 '\*' 0.05 '.' 0.1 ' ' 1

**Figure 4.**
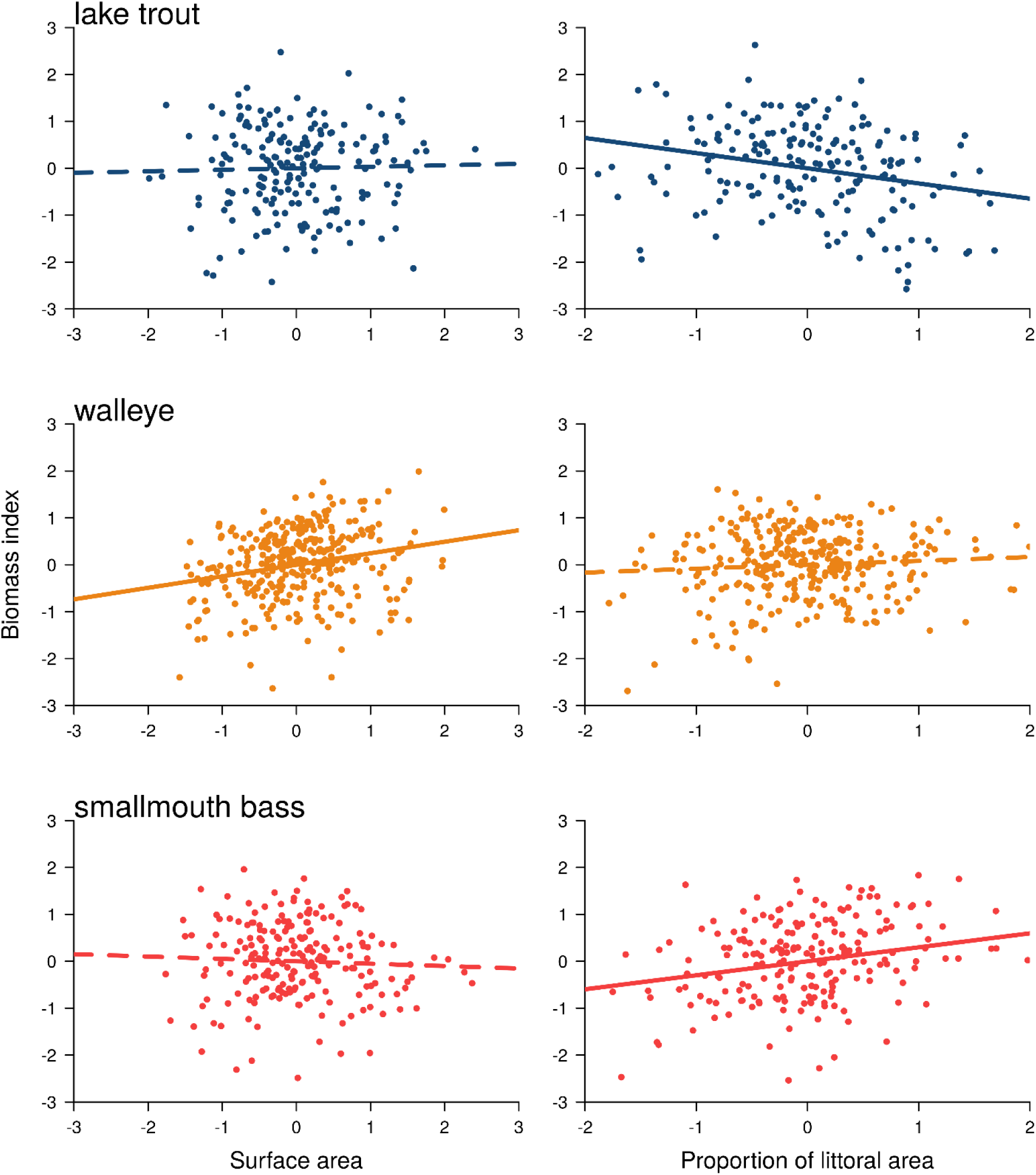
Added variable plots depicting relationships between biomass index of lake trout, walleye, and smallmouth bass with lake surface area and proportion of littoral area in Ontario lakes.

For walleye, the top-ranked model accounted for 40% of the variation in biomass index among lakes and contained the predictor variables surface area, lake maximum depth, growing degree days, shoreline development index, Secchi depth, and proportion littoral area (Table 3). Scaled coefficients indicated that growing degree days, and lake maximum depth and shoreline development index had a strong and moderate negative effect on biomass index, while lake surface area had a weak positive effect (Table 3, Fig. 2, 4).

For smallmouth bass, the top-ranked model accounted for 19% of the variation in biomass index among lakes and contained the predictor variables lake maximum depth, shoreline development index, and proportion littoral area (Table 3). Scaled coefficients indicated that lake maximum depth and shoreline development index had weak negative effects, while the proportion littoral area had a moderate positive effect (Table 3, Fig. 2, 4).

### Prey fish distribution

The top-ranked model accounted for 39% of the variation in the proportion of the prey fish captured in the offshore among lakes and contained the predictor variables lake surface area, lake maximum depth, growing degree days, and proportion littoral area (Table 4). Scaled coefficients indicated that each of these variables except surface area had a negative effect on the proportion of prey fish located in the offshore. The strongest influences on the proportion of prey fish located in the offshore were proportion of littoral area, which had a negative effect, and lake surface area, which had a positive effect on the offshore prey fish (Table 4).

**Table 4.** Results of the stepwise model selection of multiple regression testing for the influence of physical lake characteristics on the proportion of the total prey fish located in the offshore zone of each lake. SE is standard error.

| Variable | Scaled coefficient | SE | t-value | Significance<sup>1</sup> | df | F-value | R<sup>2</sup><sub>adj</sub> |
| --- | --- | --- | --- | --- | --- | --- | --- |
| intercept | 0.000 | 0.033 | 0.000 |  | 4, 549 | 90.196 | 0.392 |
| surface area | 0.323 | 0.037 | 8.740 | *** |  |  |  |
| growing degree days | -0.204 | 0.034 | -6.094 | *** |  |  |  |
| maximum depth | -0.148 | 0.048 | -3.083 | ** |  |  |  |
| proportion littoral area | -0.589 | 0.045 | -13.229 | *** |  |  |  |

<sup>1</sup>P-value significance: 0 '\*\*\*' 0.001 '\*\*' 0.01 '\*' 0.05 '.' 0.1 ' ' 1

## DISCUSSION

Predator’s responses to environmental conditions are now widely recognized as a key part of food web architecture that influences ecosystem structure and stability (Bartley et al. 2019). In this study, we sought to understand how lake morphometry impacts the depth use and biomass of three key fish predators that strongly influence food web structure while controlling for other factors, such as climate and productivity. By examining three key predator species that differ in thermal guild, we can better understand the drivers of species habitat use and related consequences for populations and for food web dynamics. Such insights are likely to be key to piecing together the fundamental factors that structure food webs, which form the backdrop for how ecosystems respond to environmental change.

We found strong evidence that species’ thermal preferences influence their responses to lake size. Consistent with our predictions, the depth use of lake trout and walleye are strongly and moderately influenced by lake surface area, respectively, while smallmouth bass show no such response in depth use. These results, therefore, support the thermal accessibility hypothesis (Dolson et al. 2009, Tunney et al. 2012), with the cold-adapted lake trout driven to occupy cold, offshore waters when nearshore waters are warm and the species is not spatially constrained in small ecosystems. Our results for lake trout also strongly resonate with previous results, which show that lake trout diet predictably vary across a range of lake size (McCann et al. 2005, Tunney et al. 2012) and that lake trout depth use is strongly shaped by temperature (Plumb and Blanchfield 2009, Guzzo et al. 2017). The moderate response of walleye may indicate that their nearshore habitat use may be limited by thermal accessibility only when temperatures in this region get high enough, allowing them to use this region more than lake trout. In contrast, and consistent with their physiology, smallmouth bass appear to have no limitations to shallow water accessibility; however, it remains unclear whether warm-water species experience limited access to the offshore. Offshore resources can be significant for some nearshore-associated predators like northern pike (*Esox lucius*) (Kennedy et al. 2018) and walleye (*Sander vitreus*) (Kaufman et al. 2009). Overall, our results are also consistent with a growing body of theory that suggests that increasing ecosystem area may impact the habitat use of predators in ways that and structure food webs in space (Eloranta et al. 2015).

We found that the thermal preferences of these predator species influenced their depth-use responses to the proportion of littoral habitat in lakes. All species occupy shallower depths as the proportion of littoral area in lakes increases, but this was stronger for walleye and smallmouth bass than for lake trout. Because of the correlation between proportion littoral area, mean lake depth, total phosphorous, and dissolved organic carbon, a larger proportion littoral area likely generally indicates a shallower, more productive lake. Thus, it seems likely that because walleye and smallmouth are less limited by thermal accessibility, these two species can respond to this increased nearshore productivity by using this area of the lake more. In contrast, the thermal limitations of lake trout (Evans 2007) limit their ability to take advantage of greater nearshore resource availability (Guzzo et al. 2017). Understanding the role of productivity and littoral area will be critical moving forward as human impacts alter these factors in boreal lakes through climate change and alteration of nearshore habitats via shoreline development and resource extraction (Keller 2007, Schindler and Lee 2010).

We did not find strong evidence that lake shape influences the depth use of top predator fishes in boreal lakes of Ontario. We found only some evidence for an influence of lake shape, estimated by shoreline development index, on the depth use of smallmouth bass but not for lake trout or walleye. This result is in contrast to previous work that shows a response of lake trout diet to changes in lake shape (Dolson et al. 2009). This discrepancy is potentially due to the vast differences in the ranges of lake area and other factors between our datasets since the lakes studied by (Dolson et al. 2009) are relatively small and geographically constrained to Algonquin Park in Ontario, Canada. It is possible that across a large geographic area, other abiotic factors are more important in driving the depth use of these species. It is also possible that lake shape does not present as much of a limitation to habitat accessibility as previously thought, especially when controlling for other factors such as littoral area and lake size. The relatively high R^2^ for lake trout depth of capture suggests that this species’ habitat use is better explained by the abiotic factors we examined here, including lake morphometry. For walleye and smallmouth bass, a significant portion of the overall variance is not explained by the models, suggesting that factors not considered in the present study could more strongly influence these species’ habitat use.

Interestingly, we found that maximum lake depth was an important factor for all species depth use, albeit in different directions. Lake trout used deeper waters in deeper lakes, while walleye and smallmouth bass both used shallower water as lakes became deeper. These asymmetric responses suggest that vertical habitat partitioning among fish of differing thermal guilds may be greater in deeper lakes (Hamrin 2008). This likely has strong implications for interactions between these species, with shallower lakes potentially forcing stronger resource competition between species of differing thermal guilds (McCann et al. 2005). These inter-guild species interactions may also be dependent on other related factors, such as the presence of preferred prey species. For example, the top-down suppression of littoral prey fish by smallmouth bass is thought to drive lake trout to feed in deeper waters (Vander Zanden et al. 1999), though this may depend on the presence of cisco (Vander Zanden et al. 2004). Thus, both vertical and horizontal spatial axes may play important roles in shaping habitat use and food web structure in these lakes (Guzzo et al. 2016).

Despite the strong relationship between depth use and lake morphometry, we find mixed support for our predictions for species’ biomass indices. Lake trout and smallmouth bass show no significant change in biomass index with lake surface area, but decrease and increase with greater proportion littoral area, respectively. In contrast, walleye shows a significant increase in biomass with lake surface area but no change with proportion littoral area. Thus, our results partially support the theoretical prediction that habitat coupling influences the biomass index of predators (McCann et al. 2005, McCauley et al. 2018). There are several possible explanations for why biomass index does not respond as clearly as depth use to lake surface area. Despite lake trout being caught deeper with increasing surface area, lake trout may not be coupling less or any change in coupling exhibited by lake trout is insufficient to influence biomass. Also, the proportion of prey captured in the offshore increases with greater lake area. This shift in prey availability may counteract changes in nearshore resource accessibility for lake trout; however, this shift in the proportion of prey in the offshore may also be a result of reduced top-down control exhibited by lake trout. In contrast, walleye biomass increased with surface area, consistent with walleye occupying deeper depths and in turn foraging on more offshore resources, such as cisco, in larger surface area lakes. Walleye are known to achieve larger lengths in lakes in which ciscoes (Kaufman et al. 2009), and larger lakes are more likely to have cisco compared to smaller lakes.

Productivity is also likely playing an important role in determining these species’ biomasses. For example, maximum lake depth showed a significant relationship with each species biomass index, likely at least in part because it correlates with productivity. Similarly, a greater proportion littoral area generally indicates a shallower, more productive lake. We found that lake trout showed a weak biomass response to increase proportion littoral area. Although lakes with small proportion littoral area are generally smaller bowl shape lakes with easily accessible littoral zones (Dolson et al. 2009), lake trout likely cannot capitalize on this suboptimal habitat due to thermal limitations, especially in summer (Christie and Regier 1988). Walleye were also caught shallower in lakes with larger proportion littoral area. Though walleye’s intermediate thermal tolerances ought to mean that they find nearshore resources more accessible than lake trout, walleye biomass did not change with proportion littoral area, suggesting that they may be similarly thermally limited. Smallmouth bass showed increased biomass with increasing proportion littoral area. Since smallmouth bass are limited to this region of the lake, they can take advantage of more resources that may be available in a larger and potentially more productive littoral zone. The coupling behaviour of smallmouth bass is poorly understood, leading to potentially spurious predictions about how much this species couples into offshore resources. However, our results that smallmouth bass are caught shallower and have increased biomass with increasing proportion littoral area might suggest that they do couple into the offshore, but exhibit reduced coupling in shallow, productive lakes. Overall, the relationship between predator habitat use and biomass index appears to be complex and requires further investigation than what we can provide here.

Our examination of the prey fish catch data shows that the proportion of prey fish located in the offshore habitat increases with lake size. This result can be interpreted in multiple ways. One possibility is that this increased biomass of prey offshore indicates an increased availability of resources in the offshore in larger lakes (i.e., a bottom-up response). Lake trout and walleye may be responding to this change in resource density rather than habitat accessibility of the nearshore. However, this would also suggest that smallmouth bass is not responding in this manner, possibly because smallmouth bass are thermally limited in terms of utilizing the cold offshore waters. Alternatively, the increase in relative offshore prey biomass may indicate reduced top-down suppression by predators despite their shift towards deep water. Our analyses here cannot tease apart these possible mechanisms; however, understanding what is driving the habitat use of these predatory species is critical for understanding what underlies food web structure in boreal lakes.

Our results have implications for how the food webs of these lakes expand and contract with lake size (Tunney et al. 2012). Previous work has shown that boreal lake trout habitat use relates to their nearshore energy use (Guzzo et al. 2017, Bartley et al. 2019). Habitat use is one key part of fish species’ behaviour that correspond to other aspects of behaviour, such as foraging (Bartley 2017). Also, the depth and distance from shore of fish predators are strongly interrelated during stratification (see Guzzo et al. 2016). Thus, the changes in the depth use that we document may indicate dietary shifts towards the offshore for lake trout and walleye but not for smallmouth bass. This deviation of predator habitat use could imply a horizontal expansion of the food web in spatially expansive environments and could contribute to resource partitioning among species differing in thermal guild. This flexible food web structure is probably an important aspect of the adaptive capacity of lake ecosystems (McMeans et al. 2016) and adds to the growing evidence that various abiotic factors can drive predator habitat use in ways that may shape local food web structure.

The link between ecosystem size and habitat use have implications for how these systems are responding to changing environmental conditions. Our work here sets the stage for understanding the how local properties of ecosystems determine their climate change responses; climate change and other human alterations to ecosystems take place in the context of local factors like lake morphometry that, taken together, configure food web structure. Understanding how these various factors interact is critical to accurately forecast the trajectories of ecosystems across the globe in the face of relentless alterations of human activities. Our work adds to a growing literature that shows how such physical properties of ecosystems as fundamental as their size determine their functioning, from individual behaviour to food web structure to ecosystem services.

## DECLARATIONS

The authors declare no competing interests.

## DATA AVAILABILITY

Data and reproducible R code will be deposited in a publicly available repository upon acceptance.

