## Supplementary Information for "Thermal preference influences depth use but not biomass of predatory fishes in response to lake morphometry"

### S1 – PRINCIPAL COMPONENTS ANALYSIS

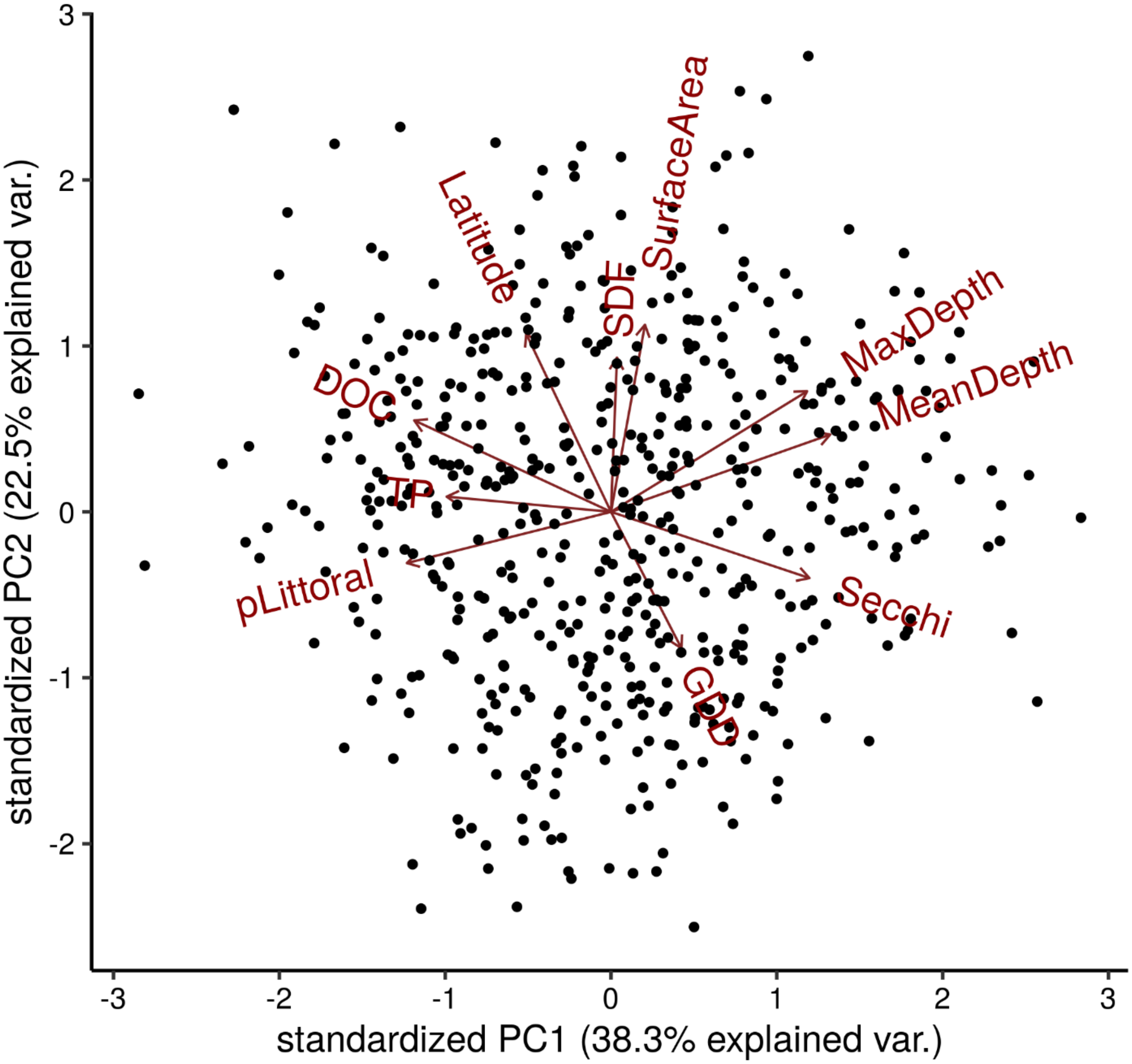


**Figure S1**. Standardized loadings of axes 1 and 2 of a principal component analysis of the transformed and scaled physical characteristics of 555 lakes included in this study. Summary statistics for these variables and details about transformation can be found in Table 1.

**Table S1**. Loadings of a principal component analysis of the transformed and scaled physical characteristics of 555 lakes included in this study. Summary statistics for these variables and details about transformation can be found in Table 1.

| Full name | Abbreviation | Comp.1 | Comp.2 | Comp.3 | Comp.4 | Comp.5 |
|---|---|---|---|---|---|---|
| surface area | SurfaceArea | | 0.491 | 0.426 | | 0.336 |
| maximum depth | MaxDepth | 0.395 | 0.316 | | -0.11 | -0.351 |
| mean depth | MeanDepth | 0.441 | 0.203 | -0.124 | -0.346 | |
| shoreline development index | SDF | | 0.405 | 0.496 | 0.389 | -0.178 |
| proportion of littoral area | pLittoral | -0.411 | -0.134 | 0.143 | 0.422 | -0.104 |
| total phosphorous | TP | -0.331 | | 0.282 | -0.554 | 0.48 |
| growing degree days | GDD | 0.142 | -0.358 | 0.542 | -0.257 | -0.118 |
| Secchi depth | Secchi | 0.4 | -0.174 | | 0.291 | 0.529 |
| latitude | Latitude | -0.171 | 0.467 | -0.394 | | 0.284 |
| dissolved organic carbon | DOC | -0.396 | 0.24 | | -0.276 | -0.328 |

**Table S2**. Summary of a principal component analysis of the transformed and scaled physical characteristics of 555 lakes included in this study. Summary statistics for these variables and details about transformation can be found in Table 1.

| **Importance of components** | **Comp.1** | **Comp.2** | **Comp.3** | **Comp.4** | **Comp.5** | **Comp.6** |
|---|---|---|---|---|---|---|
| standard deviation | 1.955045 | 1.499073 | 1.237619 | 0.961052 | 0.671406 | 0.543777 |
| proportion of variance | 0.38291 | 0.225128 | 0.153447 | 0.092529 | 0.04516 | 0.029623 |
| cumulative proportion | 0.38291 | 0.608038 | 0.761485 | 0.854013 | 0.899173 | 0.928796 |

### S2 – SUBSET ANALYSIS

**Table S3.** Results of the stepwise model selection of multiple regression testing for the influence of physical lake characteristics on the mean depth of capture of lake trout, walleye, and smallmouth bass for only lakes sampled in depth strata 1 through 5 (from 1m to 35m). SE is standard error.

***Lake trout***

| Variable | Estimate | Std. SE | t value | Pr(>\|t\|) |
|---|---|---|---|---|
| (Intercept) | 0.0000000 | 0.1148259 | 0.000000 | 1.0000000 |
| surface area | 0.4710398 | 0.1525242 | 3.088295 | 0.0033743 |
| shoreline development index | -0.2103954 | 0.1487701 | -1.414232 | 0.1638844 |
| proportion littoral area | -0.3161093 | 0.1315550 | -2.402868 | 0.0202679 |

F-statistic: 9.11888066628912 on 3 and 47 DF Adjusted R^2: 0.327565115154587

***Walleye***

| Variable | Estimate | SE | t value | Pr(>\|t\|) |
|---|---|---|---|---|
| (Intercept) | 0.0000000 | 0.0917848 | 0.000000 | 1.0000000 |
| surface area | 0.3721977 | 0.0925080 | 4.023411 | 0.0001210 |
| growing degree days | -0.1789736 | 0.0928434 | -1.927693 | 0.0571186 |
| maximum depth | -0.1569477 | 0.0961131 | -1.632947 | 0.1060534 |
| proportion littoral area | -0.2987109 | 0.0958168 | -3.117521 | 0.0024642 |

F-statistic: 7.35645837948149 on 4 and 88 DF Adjusted R^2: 0.216526745062827

***Smallmouth Bass***

| Variable | Estimate | SE | t value | Pr(>\|t\|) |
|---|---|---|---|---|
| (Intercept) | 0.0000000 | 0.1249012 | 0.000000 | 1.0000000 |
| growing degree days | 0.2414165 | 0.1315278 | 1.835479 | 0.0719440 |
| proportion littoral area | -0.3675845 | 0.1315278 | -2.794729 | 0.0071761 |

F-statistic: 4.48833578780037 on 2 and 54 DF Adjusted R^2: 0.110781840339491

**Table S4.** Results of the stepwise model selection of multiple regression testing for the influence of physical lake characteristics on the biomass index of lake trout, walleye, and smallmouth bass for only lakes sampled in depth strata 1 through 5 (from 1m to 35m). SE is standard error.

***Lake trout***

| **Variable** | **Estimate** | **SE** | **t value** | **Pr(>\|t\|)** |
|---|---|---|---|---|
| (Intercept) | 0.0000000 | 0.1292548 | 0.000000 | 1.0000000 |
| growing degree days | -0.4109118 | 0.1347194 | -3.050131 | 0.0037522 |
| Secchi depth | -0.2046783 | 0.1374099 | -1.489545 | 0.1430243 |
| proportion littoral area | -0.2384927 | 0.1341087 | -1.778353 | 0.0818148 |

F-statistic: 3.89408495886795 on 3 and 47 DF Adjusted R^2: 0.147953668359554

***Walleye***

| **Variable** | **Estimate** | **SE** | **t value** | **Pr(>\|t\|)** |
|---|---|---|---|---|
| (Intercept) | 0.0000000 | 0.0849853 | 0.000000 | 1.0000000 |
| surface area | 0.1984040 | 0.0860489 | 2.305713 | 0.0234777 |
| growing degree days | -0.4529528 | 0.0889570 | -5.091819 | 0.0000020 |
| maximum depth | -0.1336398 | 0.0906682 | -1.473944 | 0.1440651 |
| Secchi depth | -0.1897983 | 0.0931459 | -2.037644 | 0.0445889 |

F-statistic: 12.2418160544751 on 4 and 88 DF Adjusted R^2: 0.328306653963405

***Smallmouth Bass***

| **Variable** | **Estimate** | **SE** | **t value** | **Pr(>\|t\|)** |
|---|---|---|---|---|
| (Intercept) | 0.0000000 | 0.1241362 | 0.000000 | 1.0000000 |
| shoreline development index | -0.2767063 | 0.1299754 | -2.128913 | 0.0378390 |
| proportion littoral area | 0.3602332 | 0.1299754 | 2.771549 | 0.0076376 |

F-statistic: 4.87762488929691 on 2 and 54 DF Adjusted R^2: 0.121640959850771

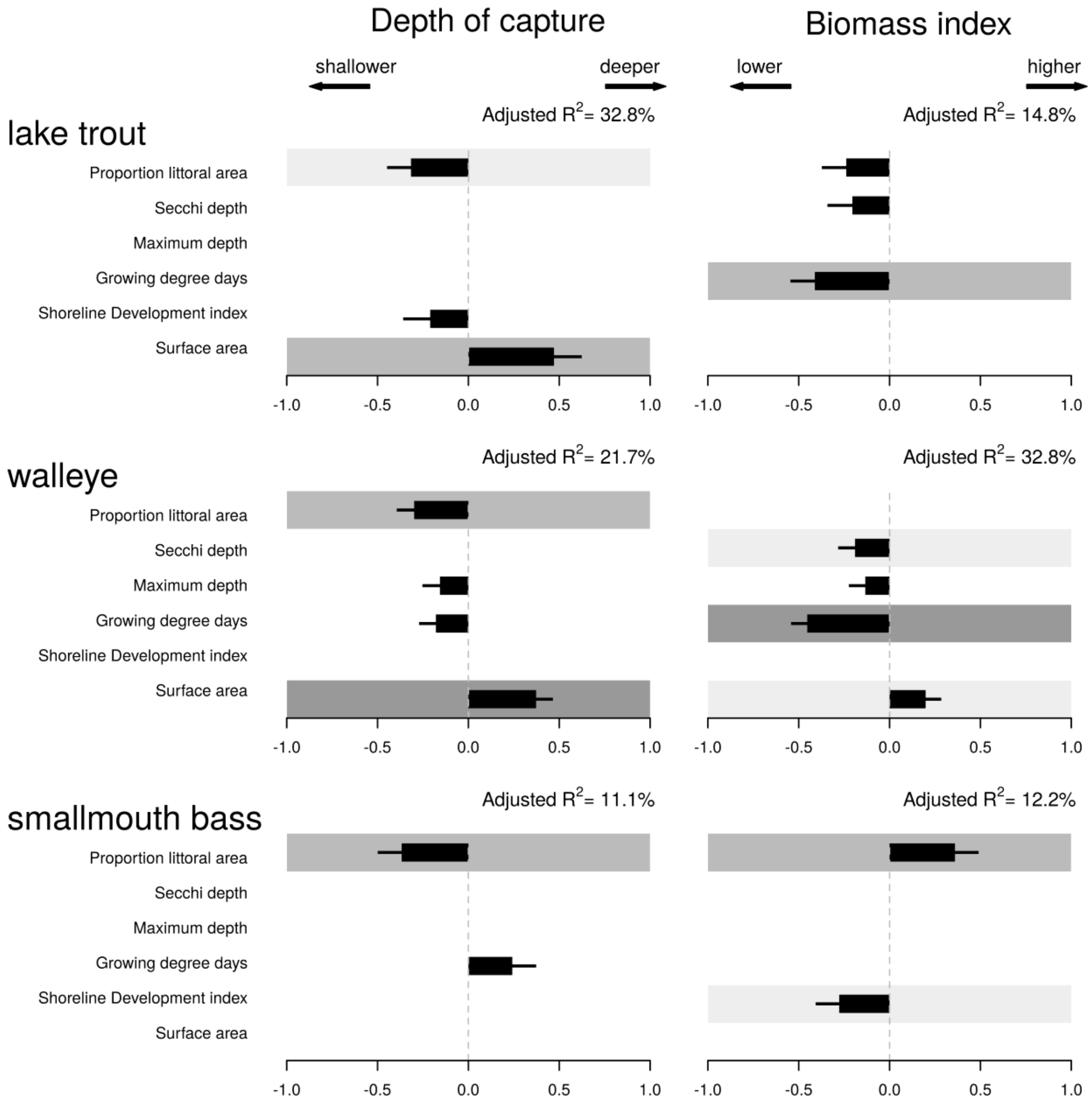


**Figure S2**. Impact of predictors on depth of capture (left side) and biomass (right side) for lake trout, walleye, and smallmouth bass for only lakes sampled in depth strata 1 through 5 (from 1m to 35m). The scaled effect of each variable included for a species and the corresponding standard error are depicted with black bars. A gray background is added to all explanatory variables found statistically significant (the different p-value thresholds 0.05, 0.01 and 0.001 are coloured in light, medium and dark gray, respectively).

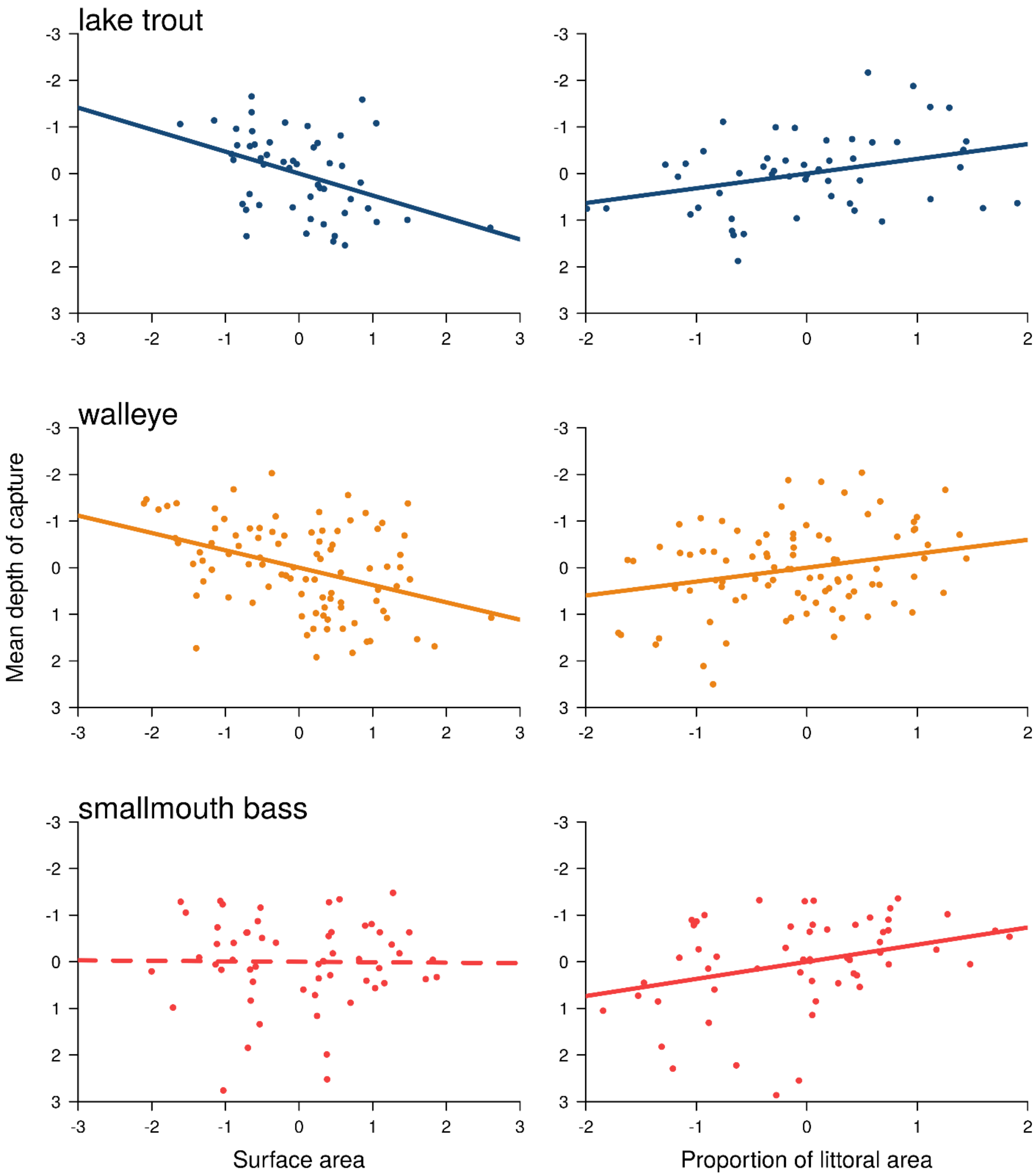


**Figure S3**. Added variable plots depicting relationships between mean depth of capture of lake trout, walleye, and smallmouth bass with lake surface area and proportion of littoral area for only lakes sampled in depth strata 1 through 5 (from 1m to 35m).

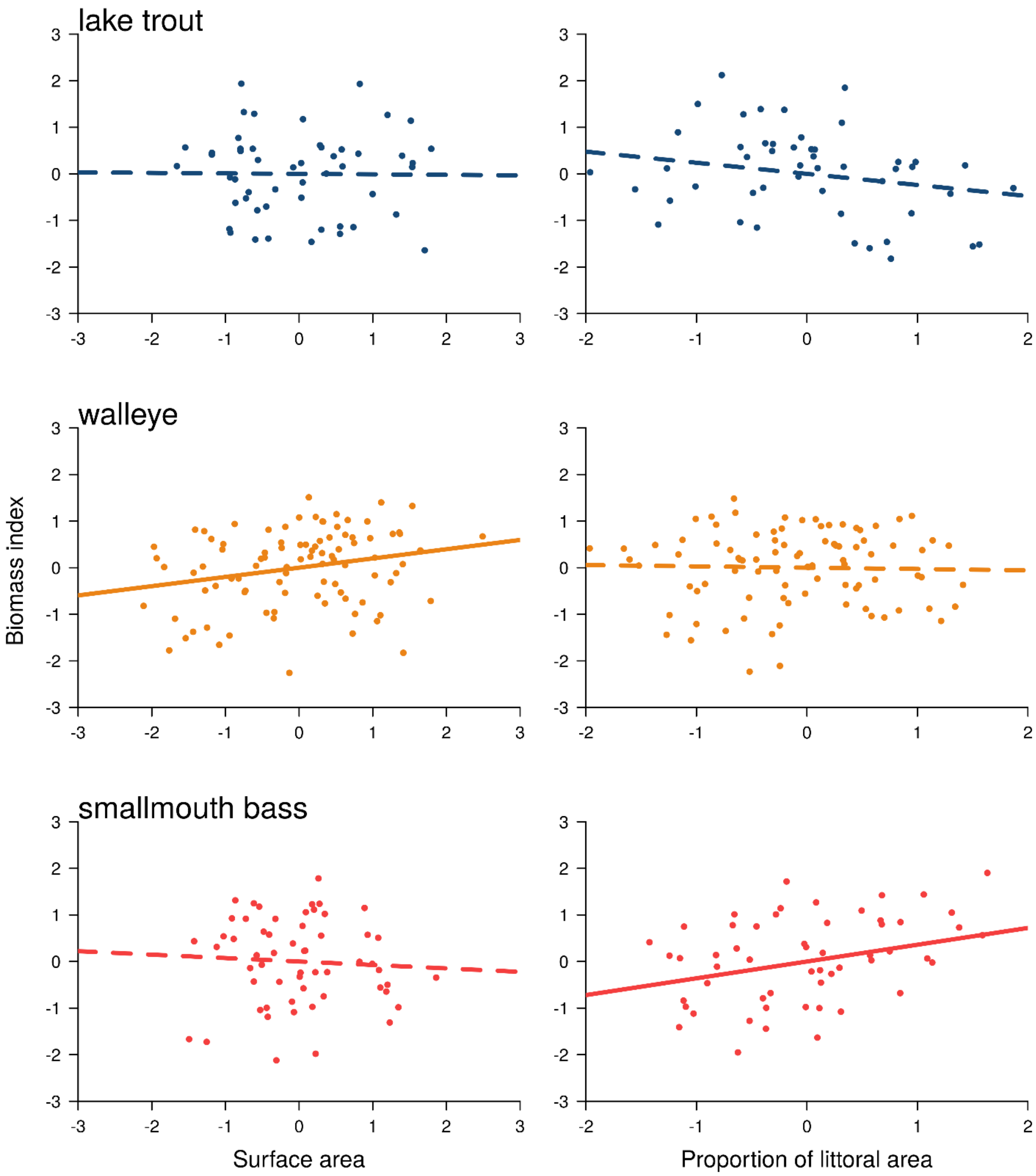


**Figure S4**. Added variable plots depicting relationships between biomass index of lake trout, walleye, and smallmouth bass with lake surface area and proportion of littoral area for only lakes sampled in depth strata 1 through 5 (from 1m to 35m).

### S3 – ANALYSIS OF CATCH-PER-UNIT-EFFORT (CUE) AND MEAN WEIGHT

**Table S5.** Results of the stepwise model selection of multiple regression testing for the influence of physical lake characteristics on catch-per-unit-effort (CUE; number per 100m nights, left side) of lake trout, walleye, and smallmouth bass. SE is standard error.

***Lake trout***

| Variable | Estimate | SE | t value | Pr(>\|t\|) |
|---|---|---|---|---|
| (Intercept) | 0.0000000 | 0.0656180 | 0.000000 | 1.0000000 |
| surface area | -0.3454316 | 0.0851343 | -4.057488 | 0.0000731 |
| growing degree days | -0.1184267 | 0.0670018 | -1.767517 | 0.0787903 |
| maximum depth | -0.2009148 | 0.0974005 | -2.062770 | 0.0405316 |
| proportion littoral area | -0.2304507 | 0.0825286 | -2.792372 | 0.0057822 |

F-statistic: 11.5067506667865 on 4 and 185 DF Adjusted R^2: 0.18191381172744

***Walleye***

| Variable | Estimate | SE | t value | Pr(>\|t\|) |
|---|---|---|---|---|
| (Intercept) | 0.0000000 | 0.0409391 | 0.000000 | 1.0000000 |
| surface area | 0.3504612 | 0.0580302 | 6.039294 | 0.0000000 |
| shoreline development index | -0.0854359 | 0.0538539 | -1.586439 | 0.1136306 |
| growing degree days | -0.4351078 | 0.0425254 | -10.231706 | 0.0000000 |
| maximum depth | -0.4614636 | 0.0495702 | -9.309290 | 0.0000000 |
| Secchi depth | -0.0754277 | 0.0469903 | -1.605178 | 0.1094440 |

F-statistic: 55.1641482381136 on 5 and 319 DF Adjusted R^2: 0.45529807963405

***Smallmouth Bass***

| Variable | Estimate | SE | t value | Pr(>\|t\|) |
|---|---|---|---|---|
| (Intercept) | 0.0000000 | 0.0629632 | 0.000000 | 1.0000000 |
| surface area | -0.2994836 | 0.0654147 | -4.578230 | 0.0000081 |
| Secchi depth | 0.1106745 | 0.0711006 | 1.556590 | 0.1210880 |
| proportion littoral area | 0.2447062 | 0.0718899 | 3.403902 | 0.0007969 |

F-statistic: 14.3525769982888 on 3 and 208 DF Adjusted R^2: 0.159555855285275

**Table S6.** Results of the stepwise model selection of multiple regression testing for the influence of physical lake characteristics on the mean weight of lake trout, walleye, and smallmouth bass. SE is standard error.

***Lake trout***

| Variable | Estimate | SE | t value | Pr(>\|t\|) |
|---|---|---|---|---|
| (Intercept) | 0.0000000 | 0.0583205 | 0.000000 | 1.0000000 |
| surface area | 0.6434243 | 0.0809997 | 7.943537 | 0.0000000 |
| shoreline development index | -0.2135860 | 0.0808089 | -2.643099 | 0.0089196 |
| growing degree days | -0.2480323 | 0.0599884 | -4.134668 | 0.0000539 |
| Secchi depth | -0.2241886 | 0.0601036 | -3.730037 | 0.0002545 |

F-statistic: 26.864769781364 on 4 and 185 DF Adjusted R^2: 0.353755743999574

***Walleye***

| Variable | Estimate | SE | t value | Pr(>\|t\|) |
|---|---|---|---|---|
| (Intercept) | 0.0000000 | 0.0531766 | 0.000000 | 1.0000000 |
| surface area | 0.1793551 | 0.0753127 | 2.381473 | 0.0178279 |
| shoreline development index | -0.2796378 | 0.0699437 | -3.998039 | 0.0000794 |
| maximum depth | 0.1398919 | 0.0642270 | 2.178084 | 0.0301296 |
| Secchi depth | 0.1508840 | 0.0592325 | 2.547320 | 0.0113230 |

F-statistic: 8.13739632806109 on 4 and 320 DF Adjusted R^2: 0.0809803400760172

***Smallmouth Bass***

| Variable | Estimate | SE | t value | Pr(>\|t\|) |
|---|---|---|---|---|
| (Intercept) | 0.0000000 | 0.0619354 | 0.000000 | 1.0000000 |
| surface area | 0.3298523 | 0.0852730 | 3.868191 | 0.0001470 |
| shoreline development index | -0.2293057 | 0.0762305 | -3.008057 | 0.0029568 |
| growing degree days | -0.2400948 | 0.0660979 | -3.632412 | 0.0003544 |
| maximum depth | -0.1739193 | 0.0798809 | -2.177231 | 0.0305998 |
| Secchi depth | -0.2411395 | 0.0682288 | -3.534279 | 0.0005048 |

F-statistic: 10.6917418441938 on 5 and 206 DF Adjusted R^2: 0.186768481838468

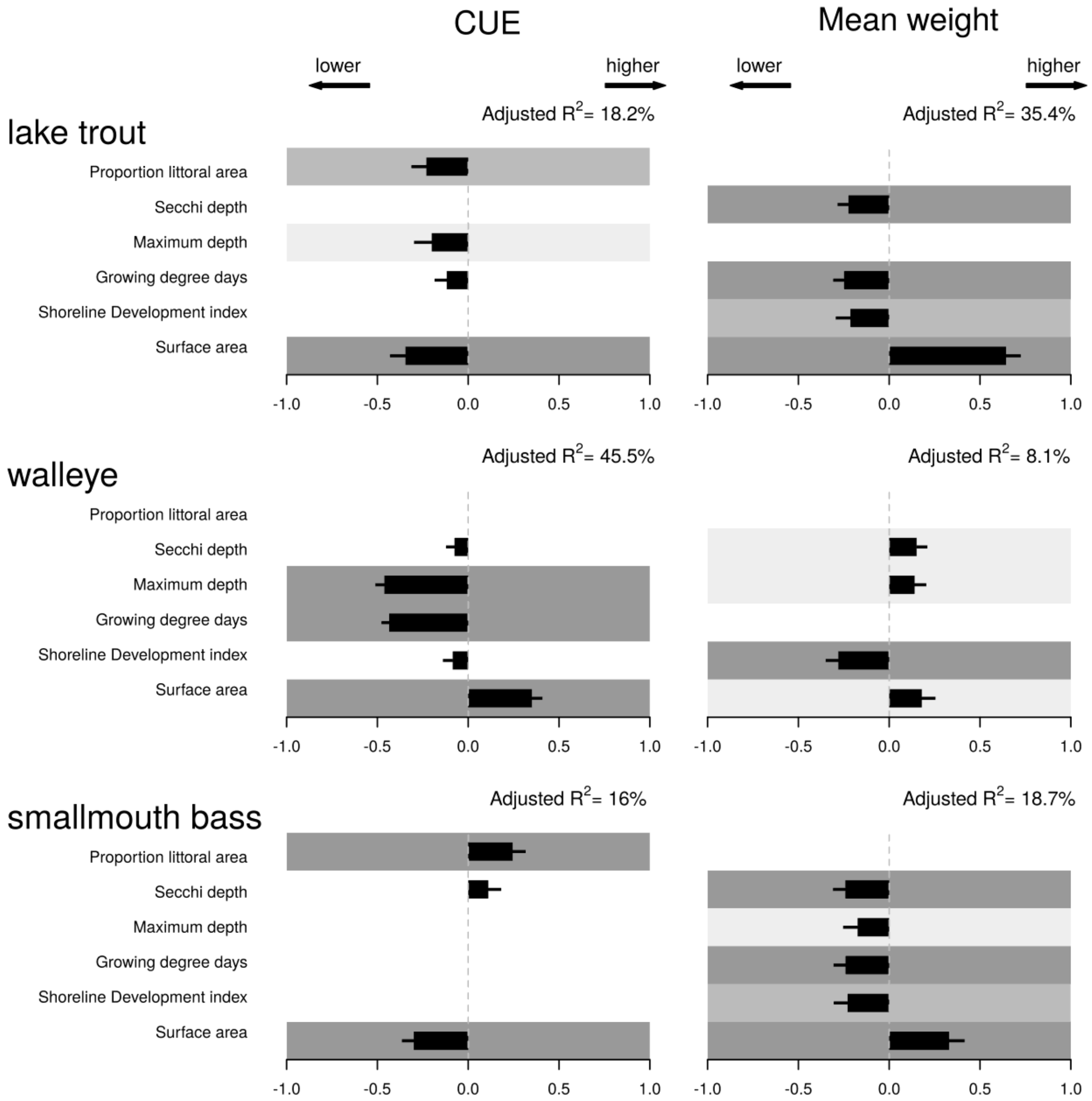


**Figure S5**. Impact of predictors on catch-per-unit-effort (CUE; number per 100m nights, left side) and mean weight (right side) for lake trout, walleye, and smallmouth bass. The scaled effect of each variable included in a species and the corresponding standard error are depicted with black bars. A gray background is added to all explanatory variables found statistically significant (the different p-value thresholds 0.05, 0.01 and 0.001 are coloured in light, medium and dark gray, respectively).

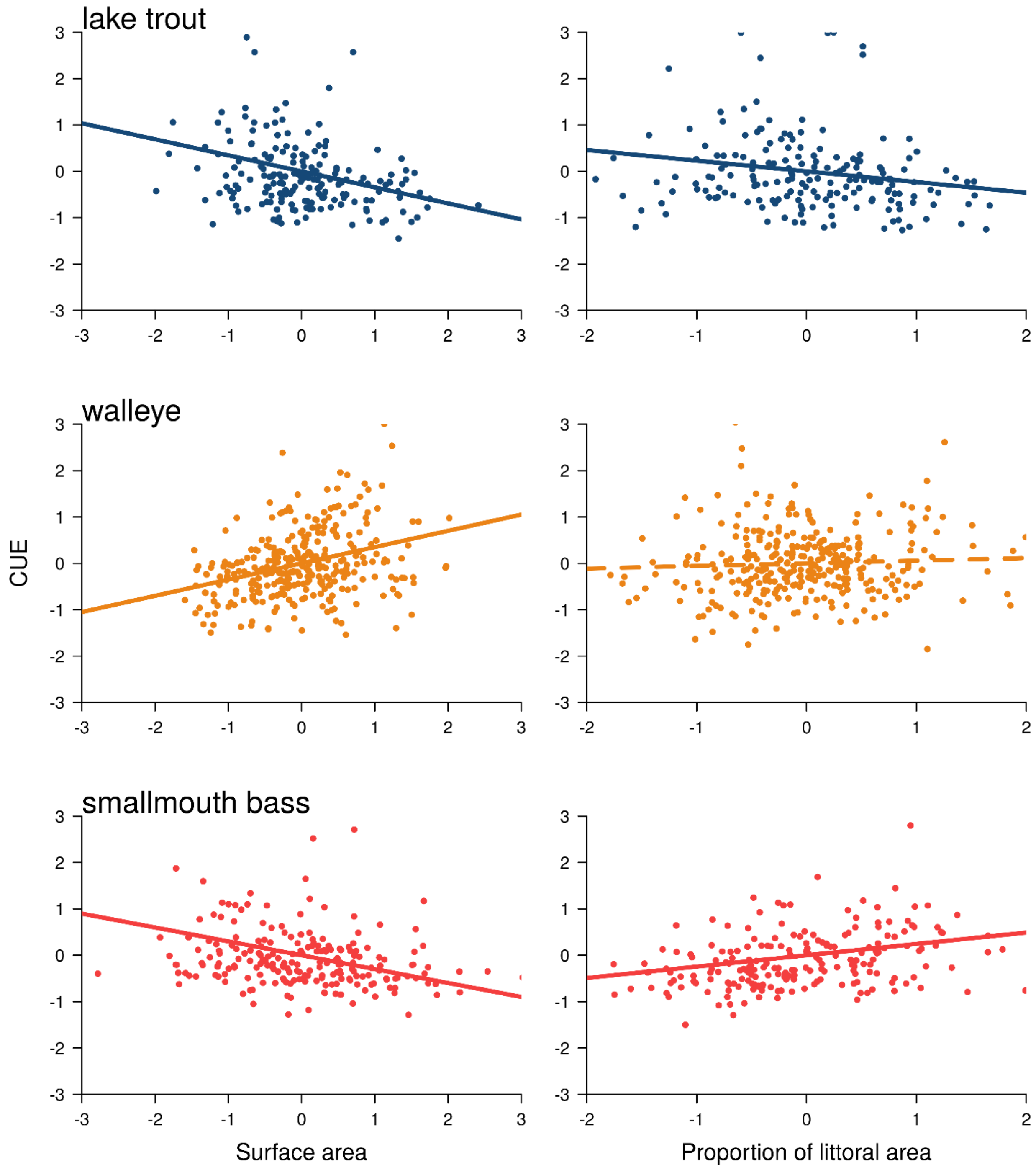


**Figure S6**. Added variable plots depicting relationships between catch-per-unit-effort (number per 100m nights) of lake trout, walleye, and smallmouth bass with lake surface area and proportion of littoral area in Ontario lakes.

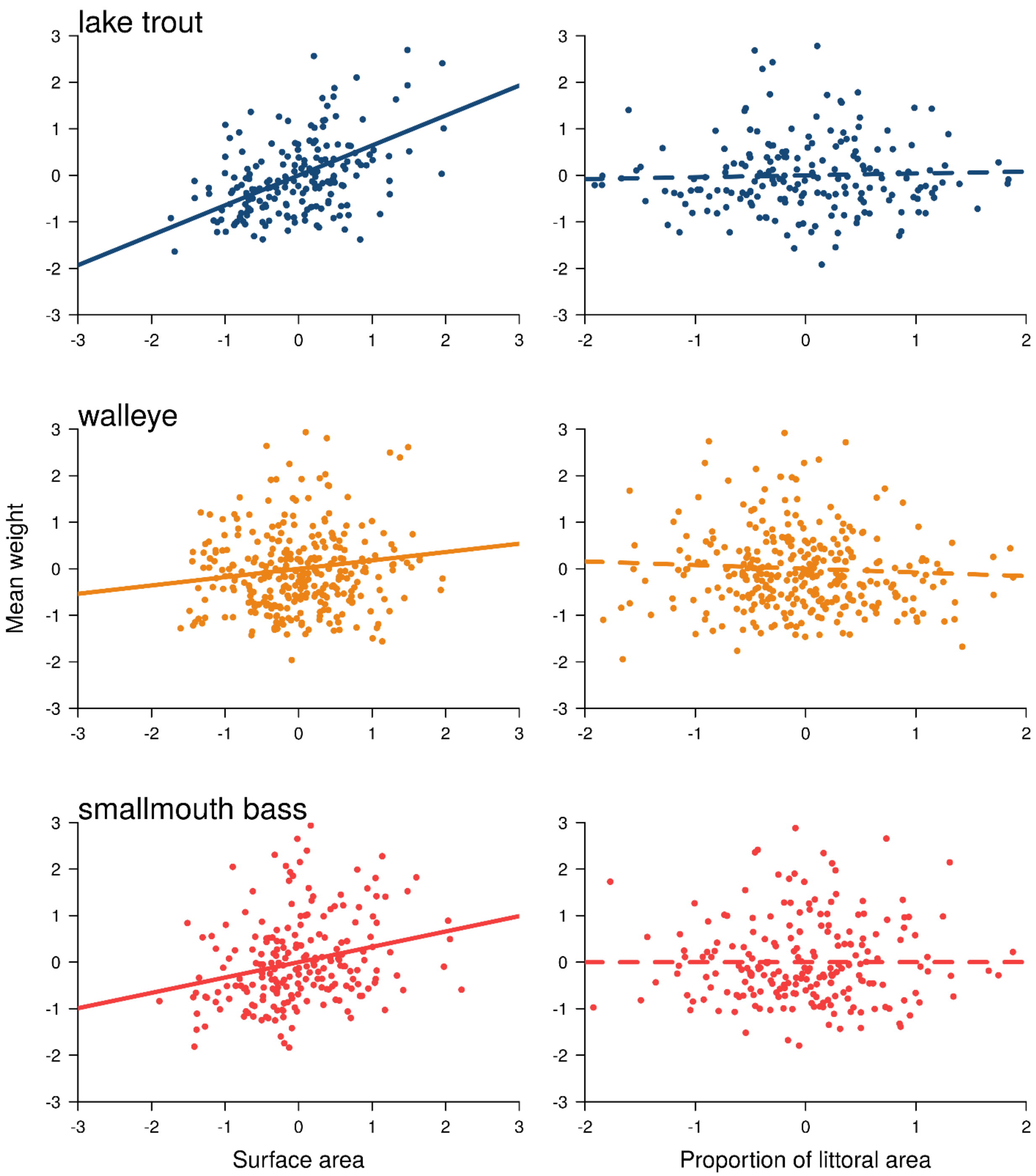


**Figure S7**. Added variable plots depicting relationships between mean weight of lake trout, walleye, and smallmouth bass with lake surface area and proportion of littoral area in Ontario lakes.
